# High-resolution structure of mammalian PI31–20S proteasome complex reveals mechanism of proteasome inhibition

**DOI:** 10.1101/2023.04.03.535455

**Authors:** Hao-Chi Hsu, Jason Wang, Abbey Kjellgren, Huilin Li, George N. DeMartino

## Abstract

Proteasome-catalyzed protein degradation mediates and regulates critical aspects of many cellular functions and is an important element of proteostasis in health and disease. Proteasome function is determined in part by the types of proteasome holoenzymes formed between the 20S core particle that catalyzes peptide bond hydrolysis and any of multiple regulatory proteins to which it binds. One of these regulators, PI31, was previously identified as an *in vitro* 20S proteasome inhibitor, but neither the molecular mechanism nor the possible physiologic significance of PI31-mediated proteasome inhibition has been clear. Here we report a high-resolution cryo-EM structure of the mammalian 20S proteasome in complex with PI31. The structure shows that two copies of the intrinsically-disordered carboxyl-terminus of PI31 are present in the central cavity of the closed-gate conformation of the proteasome and interact with proteasome catalytic sites in a manner that blocks proteolysis of substrates but resists their own degradation. The two inhibitory polypeptide chains appear to originate from PI31 monomers that enter the catalytic chamber from opposite ends of the 20S cylinder. We present evidence that PI31 can inhibit proteasome activity in mammalian cells and may serve regulatory functions for the control of cellular proteostasis.

## INTRODUCTION

Intracellular protein degradation plays a critical role in mediating and regulating many aspects of normal cellular function. Accordingly, abnormal or dysregulated protein degradation is a feature of many human diseases including cancer and those associated with muscle wasting and neurodegeneration [1–4]. Most protein degradation in eukaryotic cells is catalyzed by the proteasome, a modular protease system consisting of multiple holoenzymes composed of a common protease, the 20S proteasome, and any of various regulatory proteins including 19S/PA700, PA28αβ, PA28γ, PA200, p97 and PI31 [5, 6]. The 20S proteasome (*aka* Core Particle) is a 28-subunit protease complex composed of four axially-stacked hetero-heptameric rings [7, 8]. Each of the two identical outer rings contains seven unique α-type subunits, whereas each of the two identical inner rings contains seven unique β-type subunits, resulting in an α1-7/β1-7/β1-7/α1-7 cylindrical architecture. The β1, β2 and β5 subunits of each inner ring harbor N-terminal threonine residues that line the central interior chamber of the cylinder and function as catalytic nucleophiles for peptide bond hydrolysis with respectively different substrate specificities [7, 9]. Substrate access to these structurally-sequestered catalytic sites occurs via gated pores in the center of the α-subunit rings. Gating of the substrate access pore is an important function of regulators that compose proteasome holoenzymes. Thus, in addition to conferring specific catalytic and regulatory properties on their respective holoenzymes, such as ATP-dependent degradation of ubiquitin modified protein in the case of 19S/PA700 [10], most regulators license the complex for substrate hydrolysis by allosterically relieving constitutive pore occlusion upon binding to the apical surface of the 20S α-subunit rings [11–14].

PI31 (<u>P</u>roteasome Inhibitor of <u>31</u>,000 daltons), the product of the PSMF1 gene, was originally identified as an *in vitro* inhibitor of artificially-activated 20S proteasome [15]. PI31 has a globular N-terminus (FP domain, residues 1-151) and a natively-disordered, proline/glycine-rich C-terminus (residues 152-271) that is necessary and sufficient for proteasome inhibition (**Figure 1A**) [16]. In contrast to its well-documented *in vitro* function, the physiologic role of PI31 has been uncertain and the subject of seemingly conflicting results. For example, neither overexpression nor genetic reduction of PI31 in mammalian cells affects overall proteasome activity in a manner expected of a general proteasome inhibitor [17, 18]. Such results, as well as uncertainty about cellular roles of regulator-free 20S proteasome, suggest that PI31’s *in vitro* proteasome inhibition may lack physiologic significance. Moreover, other studies indicate that PI31 is a positive regulator of cellular proteasome function via various mechanisms whose generality and relationship to one another are not readily apparent [18–21].

**Figure 1.**
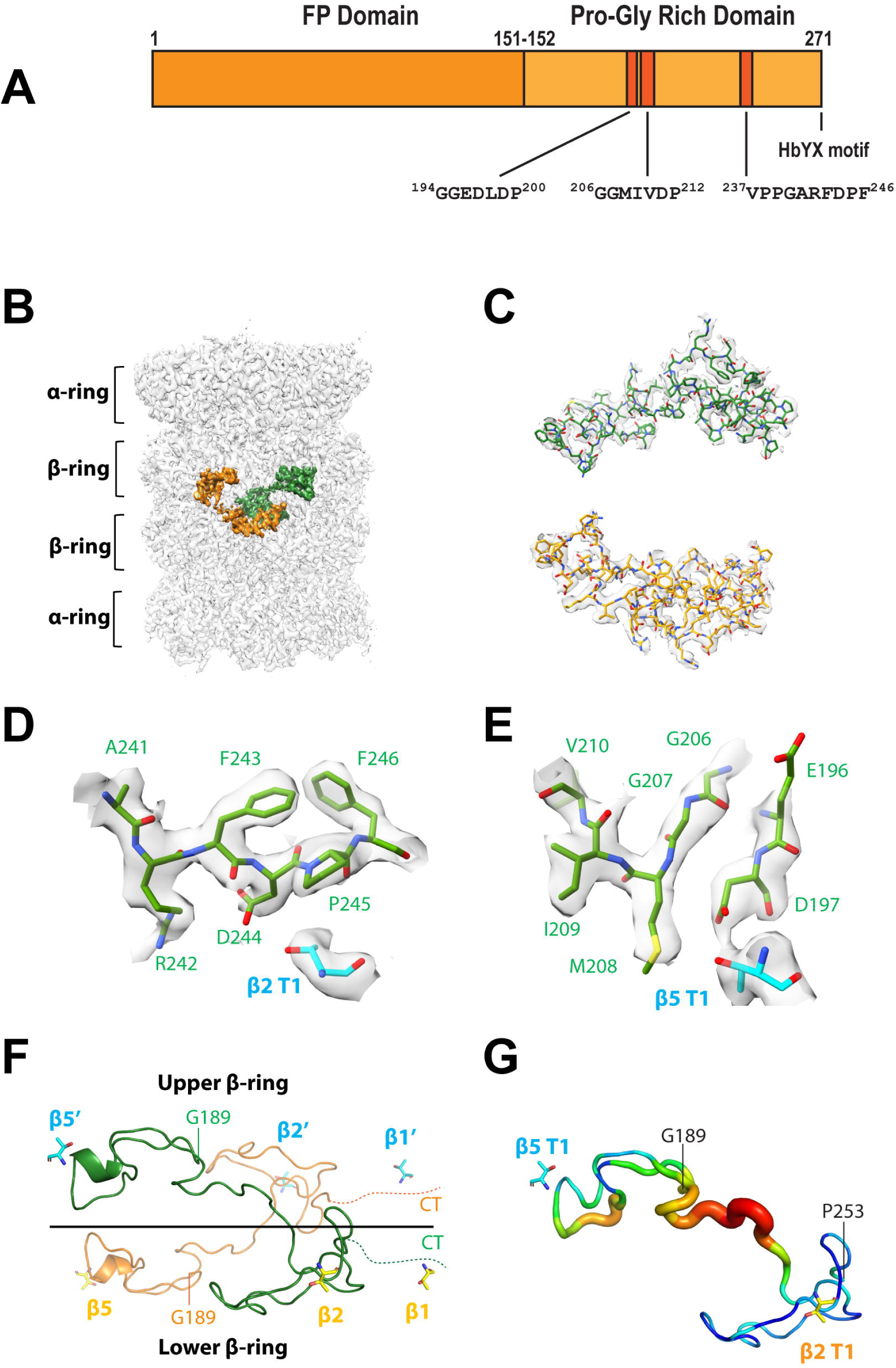
Cryo-EM structure of bovine 20S proteasome complexed with human PI31. A) Domain structure of human PI31, showing residues that interact with β5 and β2 catalytic sites (described in text). B) EM map of the human PI31-bound bovine proteasome at average resolution of 2.54 Å. The well-resolved densities belonging to the two PI31 C-terminal domians (CTDs) are shown in orange and green, respectively. C) Superposition of the PI31 atomic model with the EM density. D-E) The EM density around the β2 (D) and the β5 active site (E). F) Cartoon view of the two modeled PI31 CTDs. One PI31 CTD runs from the β5 site of one β-ring to the β2 and β1 active sites of the other β-ring. G) A B-factor plot showing that PI31 CTDs around the β2 and β5 sites are more stable. The highest and lowest B-factors are in red, and blue, respectively.

To gain further insight into physiologic roles of PI31, we solved a high resolution cryo-EM structure of a mammalian PI31-20S complex. The structure shows that the natively unstructured C-terminus of PI31 enters the central 20S catalytic chamber through the substrate access pores and interacts with catalytic sites in a manner consistent with inhibition of peptide bond hydrolysis. While our work was in progress, a comparable structure was reported for a complex between a mutant, open-gate yeast 20S proteasome and Fub1, a yeast PI31 ortholog [22]. More recently, the structure of a complex between 20S proteasome and a truncated PI31-like protein has been described from dormant spores of *Microsporidia* [23]. Although these structures are in agreement regarding a general mechanism by which PI31 interacts with and inhibits the proteasome, they have a number of specific significant differences. We have also identified conditions that manifest PI31 inhibition of proteasome function in mammalian cells and provide new context and explanations for results of earlier reports. Our work reveals new insights about the molecular and cellular roles of PI31 in regulation of proteasome function.

## RESULTS

### The C-terminal domain of human PI31 enters the central chamber of mammalian 20S proteasome and interacts directly with catalytic sites

We determined a cryo-EM structure of a mammalian PI31-20S proteasome complex after incubating recombinant human PI31 (hPI31) with the purified latent bovine 20S proteasome. The maps of PI31-free and PI31-bound proteasome were refined to the resolutions of 2.23 Å and 2.54 Å, respectively (Figure 1B, Supplemental Figures 1-2, and Supplemental Data Table 1). After model building, extra densities were identified as C-terminal domain residues (aa 189- 253) of hPI31 located in the interior of the 20S particle (Figure 1B-C). The hPI31 densities near the β2 and β5 active sites within the central chamber of 20S were especially well resolved (Figure 1D-E). Portions of two hPI31 C-terminal domains were identified within a given 20S particle and featured the same C2 symmetry characteristic of the β subunit rings. However, the C1 map was used for PI31 model building to avoid average deviations. The modeled structure is shown as a linearized hPI31 chain extending from the vicinity of the β5 active site of one β-ring and then crossing the interface of the two β-rings toward the β2 and β1 active sites of the opposing β-ring (Figure 1F). Although some hPI31 densities are present near the β1 site, we were unable to build a model at this location due to their broken nature. Because the end of the modeled hPI31 is near the β1 site, we predict an interaction between this site and the C-terminus of hPI31. Biochemical analyses described below support this prediction. The C-terminal domain of hPI31 (aa 152-271) is composed of 35% proline and glycine residues and is structurally disordered [16]. Accordingly, there is no discernable PI31 secondary structure in the resolved complex. The lack of hPI31 secondary structure indicates that hPI31 must be stabilized by the surrounding residues of the proteasome. The b-factor plot demonstrates that the hPI31 sequences near the β2 and β5 active sites are the most stable parts in the modeled hPI31 structure (Figure 1G), suggesting that the interactions between hPI31 and the proteasome active sites constitute the basis for the inhibitory activity of PI31.

Although the structure of this mammalian PI31-20S complex is generally similar to that of a recently reported complex between Fub1, the yeast PI31 ortholog, and an α3Δ yeast mutant 20S proteasome, there are a number of significant differences [22]. In each structure, two copies of the hPI31/Fub1 C-terminal domain reside within the central chamber of 20S where they appear to interact with each of the three distinct catalytic sites. Notably, the regions of hPI31/Fub1 involved in these interactions represent the most highly-conserved regions of the two proteins (Figures 2A and 2B). However, in the yeast structure, Fub1 starts (N-to-C-terminus) from the β1 active site and then extends over the center of the β-ring to the β5 active site where it dimerizes with the second copy of Fub1. The Fub1 then contacts the β5 active site of the opposing β-ring and returns to the β2 active site of original β-ring. In contrast, hPI31 does not traverse the center of proteasome in the mammalian complex. Instead, it begins from the β5 active site of one β-ring and curls toward the β2 active site of the opposing β-ring (Figures 2C and 2D). Thus, although PI31 and Fub1 have conserved active-site interacting regions, the structural arrangement of these interactions appears to be distinct (Figures 2B-2D). Whereas Fub1 blocks the β5 activity through a unique conformation between two separate Fub1 molecules, hPI31 achieves this conformation using two conserved sequences (aa 194-200 and aa 206-212) within the same chain near the β5 active site. Specifically, the antiparallel hydrogen bonding between Gly194-Gly195-Glu196 and Gly206-Gly207 creates a conformation that fits into the β5 binding site (Figure 2E). Human PI31 mimics the substrate binding, P4-P3-P2-P1- P1’-P2’-P3’-P4’ to S4-S3-S2-S1-S1’-S2’-S3’-S4’, to inhibit the proteolytic activity of proteasome. At the β2 active site, residues 242-246 of hPI31 occupy the β2 pocket in an antiparallel direction with Asp244 (P1) and Arg242 (P3) inserting into the S1 and S3 binding pockets, respectively (Figure 3A). The hPI31 residues Arg242, Asp 244, Pro245, and Phe246 are hydrogen bonded with the surrounding residues Thr1, Gly47, Ala49, Gly128, and Ser129 of β2 subunit to stabilize the binding sequence (Figure 3B). At the β5 active site, the dipeptide Asp197-Leu198 and residues 208-214 of hPI31 fit into the β5 binding pocket in a parallel direction with the Met208(P1) and Ile209(P3) inserting into the S1 and S3 binding pockets (Figure 3C). The hPI31 residues Asp197, Met208, and Ile209 are hydrogen bonded with the surrounding residues Thr1, Thr21, Gly47, Ala49, and Ser130 of β5 subunit to sustain the unique binding conformation (Figure 3D).

**Figure 2.**
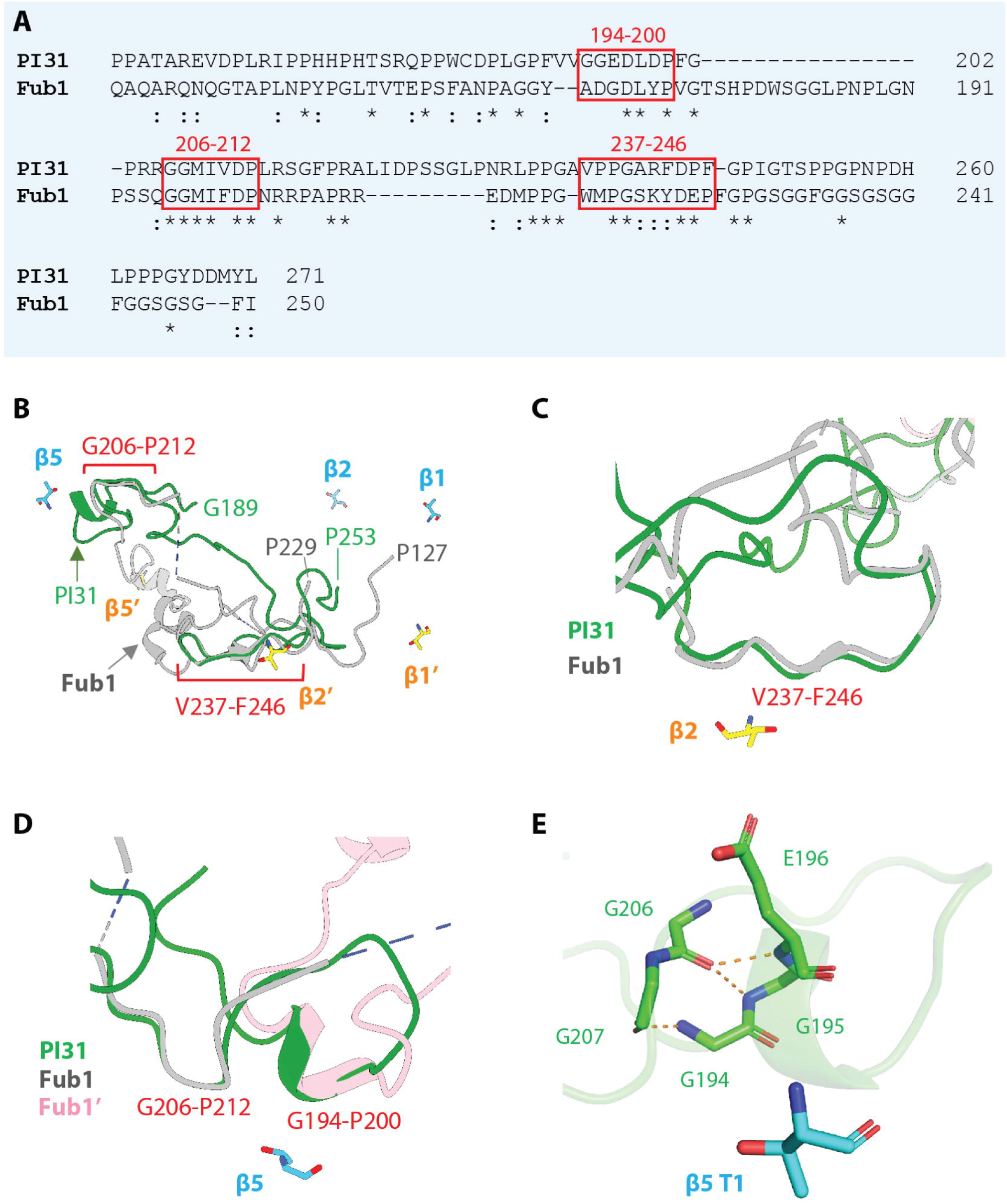
Similarities and differences between PI31 and Fub1 in 20S binding. A) Sequence alignment of the C-terminal regions of human PI31 (aa 159-271) and yeast Fub1 (aa 134-250). Structurally conserved sequences are show in red squares. B ) Structural alignment of the C-terminal regions of human PI31 and yeast Fub1. The β5 subunit of the human PI31- bovine proteasome structure was aligned with that of Fub1-α3Δ yeast proteasome mutant structure. Only the regions near the β2 and β5 active site are structurally conserved. The structurally conserved regions are shown in red squares. C) Zoomed view of the aligned structures of PI31 and Fub1 near the β2 site. D) Zoomed view of the aligned structures of PI31 and Fub1 near the β5 site. Fub1 uses two molecules of Fub1 C-terminals to occupy the β5 site. Instead, PI31 forms a unique conformation within the same molecule to block the β5 site. E) The special conformation of PI31 near β5 is constructed by the antiparallel hydrogen bonding between the main chains of the two conserved sequences, Gly194-Gly195-Glu196 and Gly206- Gly207.

**Figure 3.**
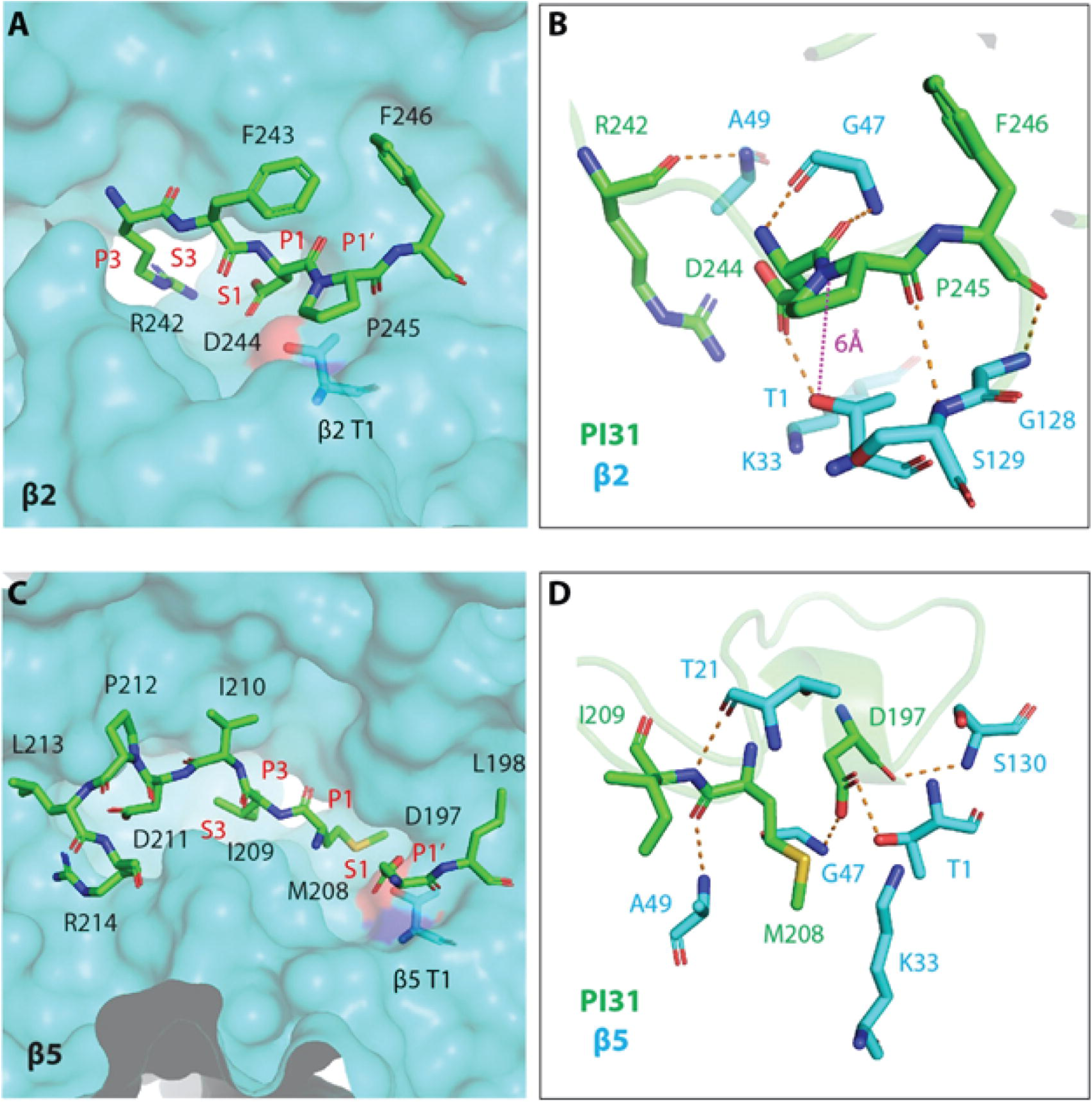
PI31 mimics substrate binding to the proteasome active sites. A) Binding of hPI31 to the β2 active site. The side chains of Asp244 (P1) and Arg242 (P3) are inserted into the S1 and S3 binding pockets, respectively. The Pro245 is at the P1’ position. B) PI31 in the β2 active site was stabilized by the surrounding amino acids of the β2 site. The distance between Thr1 and the cleavage site is 6 Å and may contribute to the proteolytic resistance to the β2 site. C) Binding of hPI31 to the β5 active site. The side chains of M208 (P1) and I209 (P3) are inserted into the S1 and S3 binding pockets, respectively. Asp197 is at the P1’ position. D) Hydrogen bonds between PI31 and the surrounding amino acids of β5 help to stabilize the bound special conformation of PI31 in the β5. The special conformation leaves no peptide bond between the P1 (Met208) and P1’ (Asp197) for the β5 activity.

### PI31 interacts with catalytic sites of the proteasome in a manner that prevents its own degradation

The ability of PI31 to stably inhibit the proteasome requires that PI31 itself is resistant to degradation. Several features at the β2 active site may contribute to this resistance. First, Lys33 of the β2 subunit was proposed to promote the deprotonation of the catalytic Thr1 and to generate the oxyanion for nucleophilic attack [24, 25]. The hydrogen bond between OD1 of Asp244 of hPI31 and HG1 of Thr1 may impede the deprotonation of HG1 from OG1 of Thr1 by Lys33 and prevent the formation of the threonine nucleophile. Second, the trypsin-like activity the β2 active site has a preference for basic residues at the P1 site. In the structure of the PI31- β2 site, P1 is occupied by Asp244 which is unfavorable for hydrolysis. According to the Keil rule, trypsin can cleave after Arg or Lys, but not at those sites before Pro [26]. Therefore, Pro245 at the P1’ site likely hampers the trypsin-like activity toward PI31 at the β2 active site. Third, the distance between the cleavage site (P1’-P1) and the catalytic Thr1 is 6 Å (Figure 3B), which further restrains the proteolytic activity of the β2 active site. At the β5 site, Asp197 of PI31 forms a hydrogen bond with the β5 catalytic Thr1 (Figure 3D). Like the hydrogen bonding between Asp244 and Thr1 of β2, the hydrogen bond between Asp197 and Thr1 of β5 may prevent the formation of threonine nucleophile at the β5 site. Additionally, the unique antiparallel conformation formed by Gly194-Gly195-Glu196 and Gly206-Gly207 of PI31 forces Asp197 to sit at the S1’ site and residues 208-214 to bind to β5 pocket in a parallel direction, which is in the reverse direction of normal substrate binding. This binding profile leaves no peptide bond between P1 (Met208) and P1’ (Asp197) and renders PI31 unfavorable for hydrolysis at the β5 site.

### The C-terminal domain of hPI31 breaches the closed gate conformation of wild-type mammalian 20S proteasome

The mammalian PI31-20S complex structure reported here was obtained with latent wild-type 20S proteasome featuring a constitutively closed gate conformation. This form of the proteasome exhibits low protease activity in the absence of endogenous gate-opening activators, raising questions about how hPI31 gains access to the structurally sequestered 20S catalytic sites. Although PI31 was present in 5-fold molar excess of 20S in samples prepared for cryo-EM analysis, only 34% of 20S particles contained extra densities after extensive 3D classification (Supplemental Figure 2). This finding suggests that structural and/or functional features of these proteins limit their interaction. It is possible that the C-terminus of PI31 enters the proteasome during transient openings of the predominately closed-gate conformation during the 20S-PI31 pre-incubation [27, 28]. Alternatively, our preparations of bovine 20S proteasome might contain low levels of constitutively activated (i.e., open-gate) 20S known to be promoted by certain purification conditions [29]. Either possibility would be consistent with the relatively low efficiency of complex formation revealed by 3D classification even after prolonged incubation of 20S with a molar excess of PI31. In either case, we predicted that hPI31 should be a much more efficient inhibitor of open-gated 20S. To test this prediction, we compared PI31 inhibition of 20S in the presence and absence of low concentrations of SDS, commonly used to enhance proteasome activity by opening the substrate access gates. In fact, when 20S was activated by low concentrations of SDS, PI31 inhibited proteasome activity to a much greater extent than with latent 20S (Figure 4). These results indicate that PI31 readily gains access to the proteasome’s catalytic sites when its C-terminus passes through the central pore of the apical α-subunit rings.

**Figure 4.**
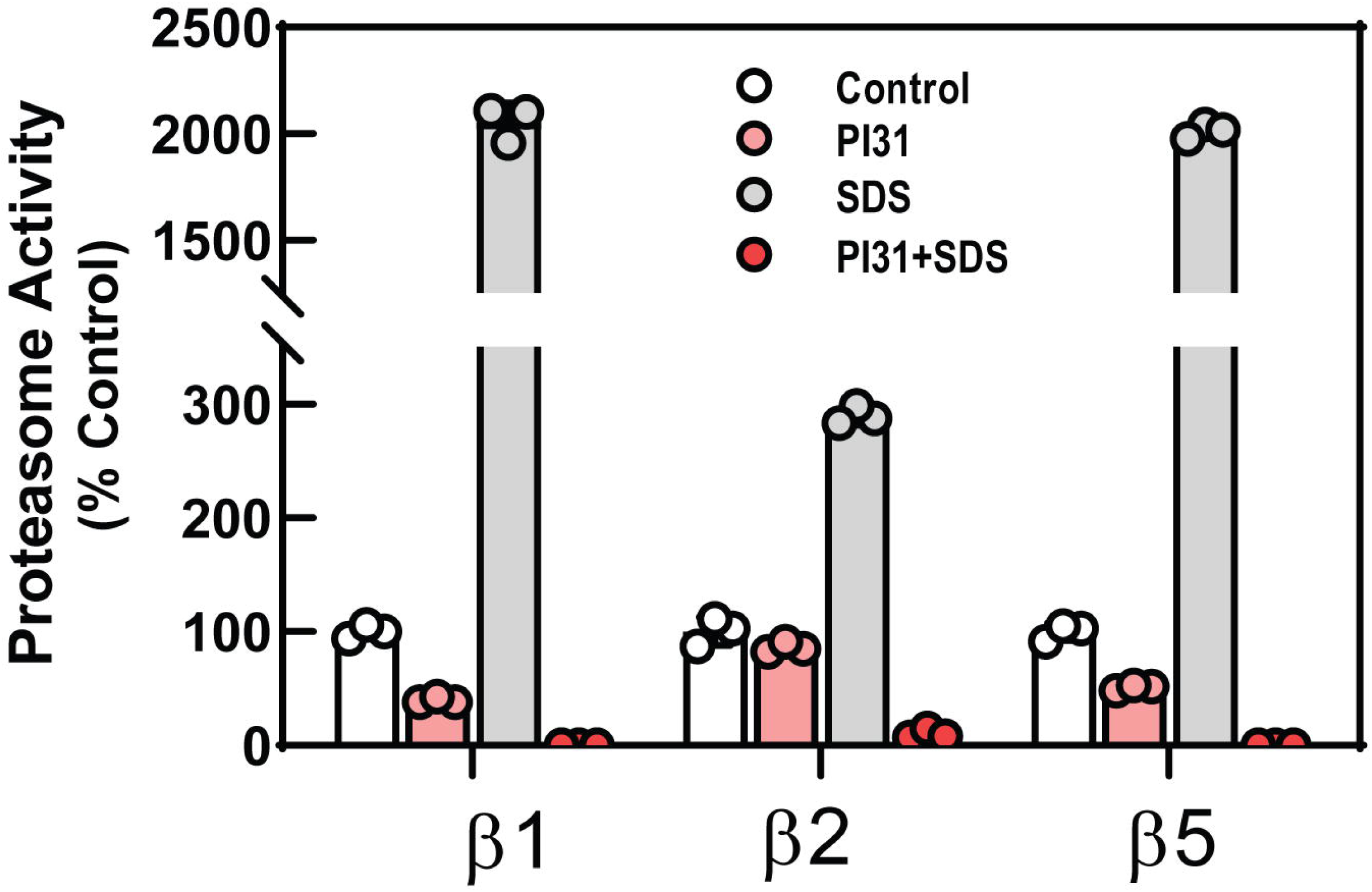
Open-gated 20S proteasome is inhibited more effectively than closed gate proteasome by PI31. Purified wild-type hPI31 was incubated with purified latent 20S proteasome in the absence or presence of 0.03% SDS prior to assay of proteasome activity with peptide substrates specific for indicated catalytic subunits, as described in Experimental Procedures. Proteasome activity in the absence of SDS or PI31 was set to 100 and all other activities are expressed as a percentage of that value. Bars represent mean values of indicated triplicate assays. Similar data were obtained in at least four independent experiments.

### Individual catalytic subunit types are inhibited by specific sequences of hPI31

The structure of the hPI31-20S complex described above predicts hPI31 residues likely to be involved in inhibition of specific catalytic sites. To test these predictions, we first expressed and purified hPI31 truncation mutants that contain or lacked various combinations of these residues and tested these mutants in activity assays specific for each of the proteasome’s three catalytic subunit types (Figure 5A). As expected, both full-length wild-type hPI31 and a truncation mutant lacking the FP domain (residues 1-151) but containing the entire C-terminal domain (residues 152-271) inhibited the catalytic activity of each subunit type by > 95% (Figure 5B). Similarly, a mutant consisting of residues 1-261 inhibited activity of each catalytic subunit type. In contrast, a mutant consisting of residues 1-181 (i.e. residues N-terminal to any observed catalytic site-interacting residues of PI31) had no inhibitory activity. These results confirm that PI31 amino acids responsible for inhibition of proteasome active sites reside between residues 182-262. Accordingly, PI31 variants 1-244 and 152-244 each inhibited β5 activity but not β1 activity, as predicted. However, neither of these latter mutants inhibited β2 activity, as expected, but instead stimulated β2-catalyzed hydrolysis. This surprising result suggests that these truncated proteins lack residues required for the stable orientation of the amino acid(s) that contact(s) the active site β2 threonine. The differential effects of these PI31 mutants on specific types of proteasome activities were also demonstrated in an orthogonal assay using the activity probe, Me_4_BodipyFL-Ahx_3_Leu_3_-VS (Figure 5C). Me_4_BodipyFL-Ahx_3_Leu_3_-VS covalently binds to each of the proteasome’s catalytically active sites and the resulting fluorescently tagged subunits can be detected and identified after SDS-PAGE [30]. Thus, inhibitors of any catalytic site will reduce labeling of that respective subunit. As expected, both full-length, wild-type and the C-terminal domain (residues 152-271) PI31s, greatly reduced labeling of each proteasome subunit type, whereas PI31^1-181^ had no effect on labeling of any subunit. In contrast, a mutant consisting of residues 1-261 inhibited labeling of each catalytic subunit type (Figure 5C). These functional results are consistent with the structural predictions for identification of interaction sites between PI31 and individual catalytic subunits that cause proteasome inhibition and indicate that despite a clear structural resolution of PI31 residues in contact with the β1 catalytic site the latter interaction occurs between PI31 residues 245-261.

**Figure 5.**
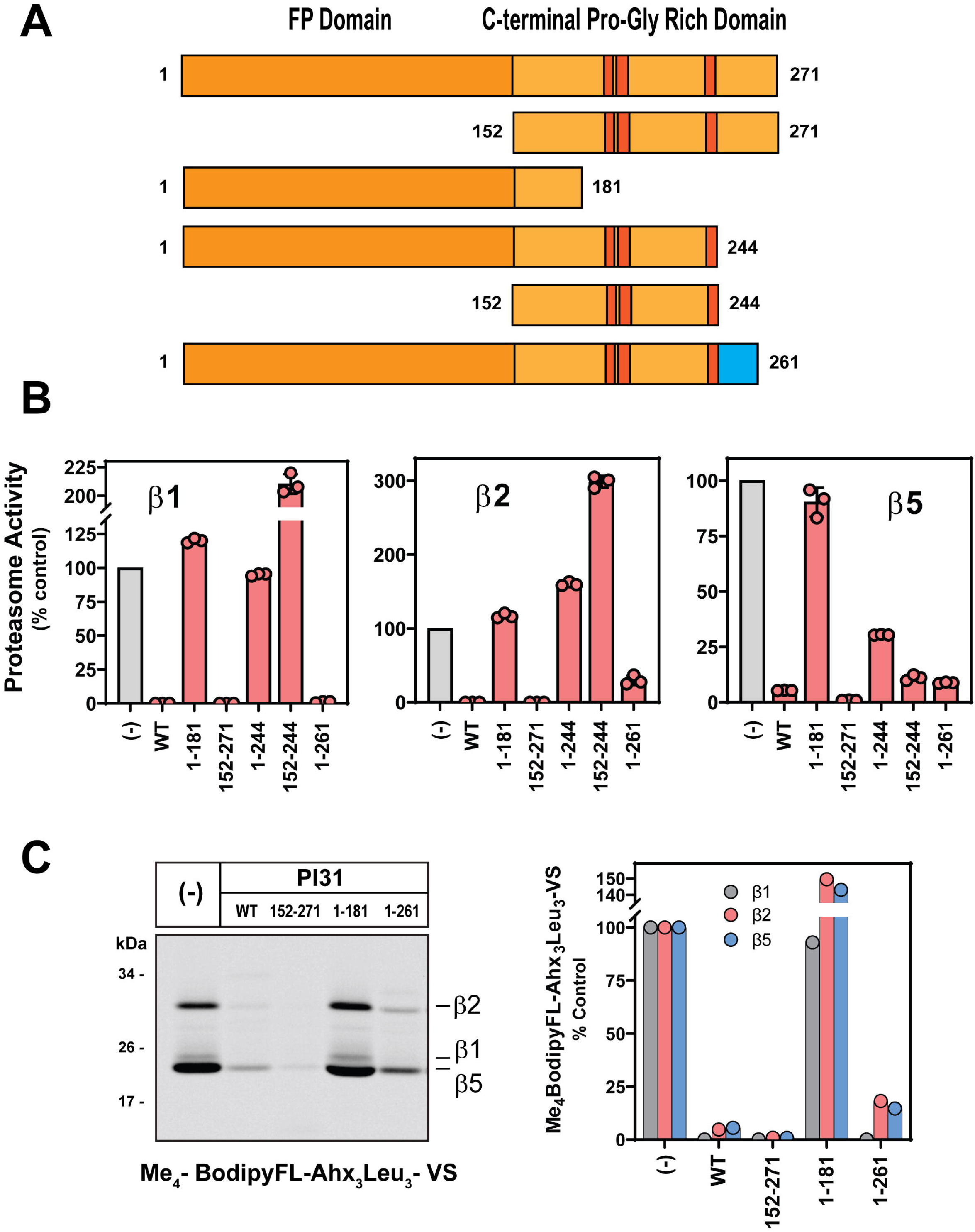
Identification of PI31 interaction sites with different 20S proteasome catalytic subunit types. A) Recombinant wild-type and indicated truncation mutants of hPI31 were expressed and purified as described in Experimental Procedures. Mutants are depicted schematically and show proposed interaction sites for β5, β2 and β1 catalytic sites (see Figure 1A). The proposed interaction site for the β1 subunit is indicated in blue. B) 20S proteasome (50 nM) activity for each catalytic subunit type was measured in the absence (-) or presence of wild-type and indicated hPI31 mutants at 10 to 20-fold molar excess. Proteasome activity for each subunit type in the absence of PI31 (-) was set to 100 and other activities were express as a percentage of that value. Bars represent mean values (± SD) of indicated triplicate assays. Similar results were obtained in three independent experiments. C) Activated 20S proteasome (115 nM) was pre-incubated in the absence (-) or presence of indicated hPI31 variants (10 µM) and then labeled with Me_4_BodipyFL-Ahx_3_Leu_3_-VS probe (500 nM) for 15 minutes to avoid signal saturation. Assays were quenched with SDS-PAGE sample buffer and samples were subjected to SDS-PAGE and imaging as described in Experimental Procedures.

To further refine the functional specificity of interactions between individual 20S catalytic sites and corresponding PI31 residues involved in inhibition, we expressed and purified hPI31 with point mutations (D to A) at resides 197 and 244. Our structure shows that these residues interacted directly with the catalytic threonine residues of β5 and β2 proteasome subunits, respectively (Figure 3). Because of the unresolved nature of the interaction between hPI31 and the β1 site, we cannot confidently predict a specific mutation that would allow an analogous test for this site. Consistent our predictions, hPI31^D197A^ inhibited β5-specific peptide hydrolysis 3- times less well than wild-type hPI31 but had little or no effect on the inhibition of hydrolysis of β1- or β2-specific substrates compared to wild-type hPI31 activities (Figure 6A). A similar β5- specific diminution PI31 inhibition by the hPI31^D197A^ mutant was observed in an assay using the active-site probe, Me_4_BodipyFL-Ahx_3_Leu_3_-VS (Figure 6B). Likewise, hPI31^D244A^ had significantly weaker inhibitory activity against β2-specific peptidase activity but did not affect inhibition of the β5-specific substrate hydrolysis (Figure 6A). This latter mutant also blocked normal PI31 inhibition of β1-specific substrate hydrolysis to a lesser extent. Although the basis of this effect is unclear, we note that the proteasome’s catalytic sites are known to have interdependent allosteric effects [31]. Therefore, selective inhibition at one or more catalytic sites may affect activity at other sites.

**Figure 6.**
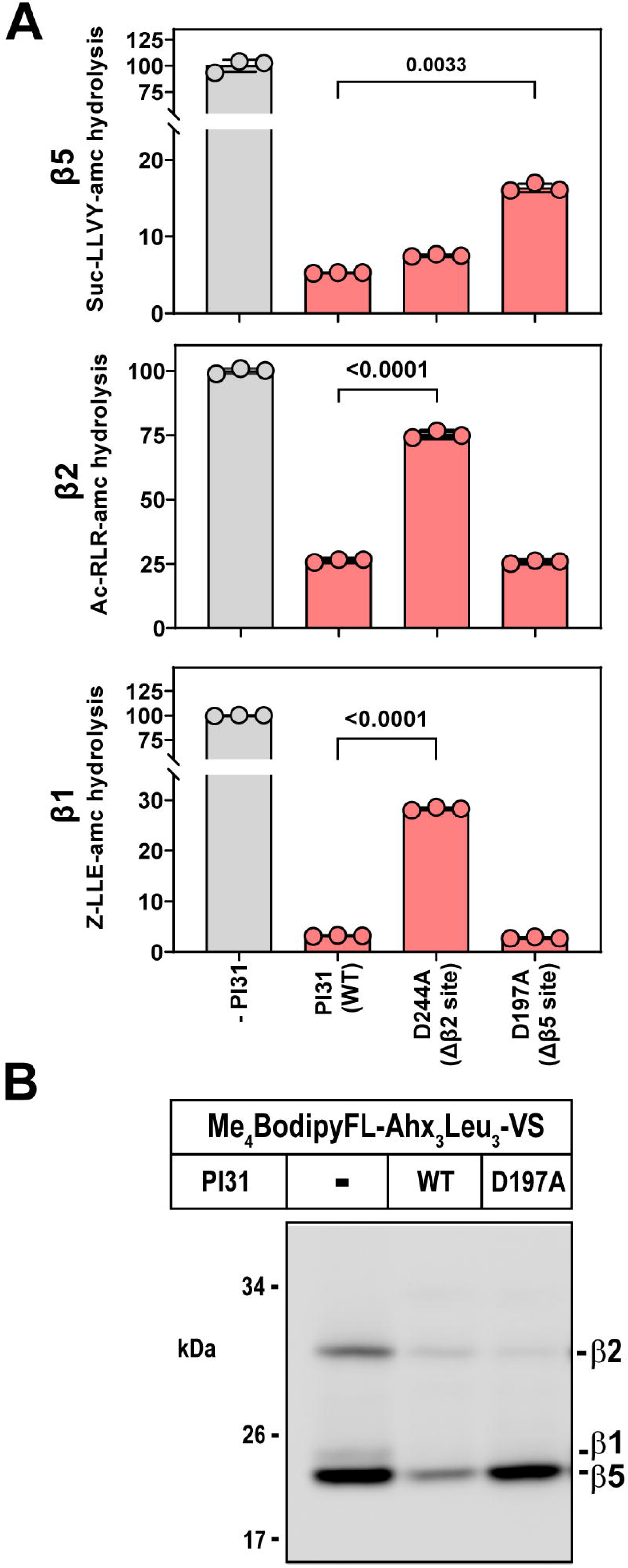
Identification of individual PI31 residues involved in site-specific inhibition of proteasome activity. A) 20S proteasome (143 nM) activity for each indicated catalytic sites was measured as described in Experimental procedures following incubation with 10-fold molar excess of PI31 (1.4 µM). Activity in the absence of PI31 was set to 100 and other activities are expressed as a percentage of that value. Bars represent mean values (± SD) of indicated triplicate assays. Similar results were obtained in three independent experiments. Differences in proteasome activity were analyzed by one-way ANOVA and Tukey’s HSD posthoc test. P-values are shown for significant differences between PI31 treatments. B) 20S proteasome (143 nM) was activated by SDS and incubated in the absence or presence of indicated variants of PI31 (1.4 µM). Samples were then labeled with Me_4_BodipyFL-Ahx_3_Leu_3_-VS activity-based probe (1 µM) for 40 minutes. Assays were quenched with SDS-PAGE sample buffer and samples were subjected to SDS-PAGE and imaging as described in Experimental Procedures.

### Monomeric PI31 is sufficient for proteasome inhibition

Previous studies established that PI31 forms homodimers mediated by the N-terminal FP domain [16, 32]. However, the ability of the isolated C-terminal domain to fully inhibit proteasome activity suggests that monomeric PI31 is the functional form of the protein. To test this hypothesis further, we expressed and purified hPI31 containing a point mutation (V6R) in the FP domain predicted to disrupt PI31 dimerization [32]. We used mass photometry to confirm that wild type hPI31 exists in a monomer-dimer equilibrium at concentrations likely to reflect those of cellular conditions (Figure 7A). The V6R mutation significantly shifted the monomer-dimer equilibrium in favor of monomeric PI31 which retained inhibitory activity against proteasome catalysis (Figure 7B and C). These functional results are most reasonably explained by a structural model in which the two molecules of PI31 observed in the catalytic chamber of 20S originate from two PI31 monomers that enter from opposite ends of the 20S cylinder (Figure 8).

**Figure 7.**
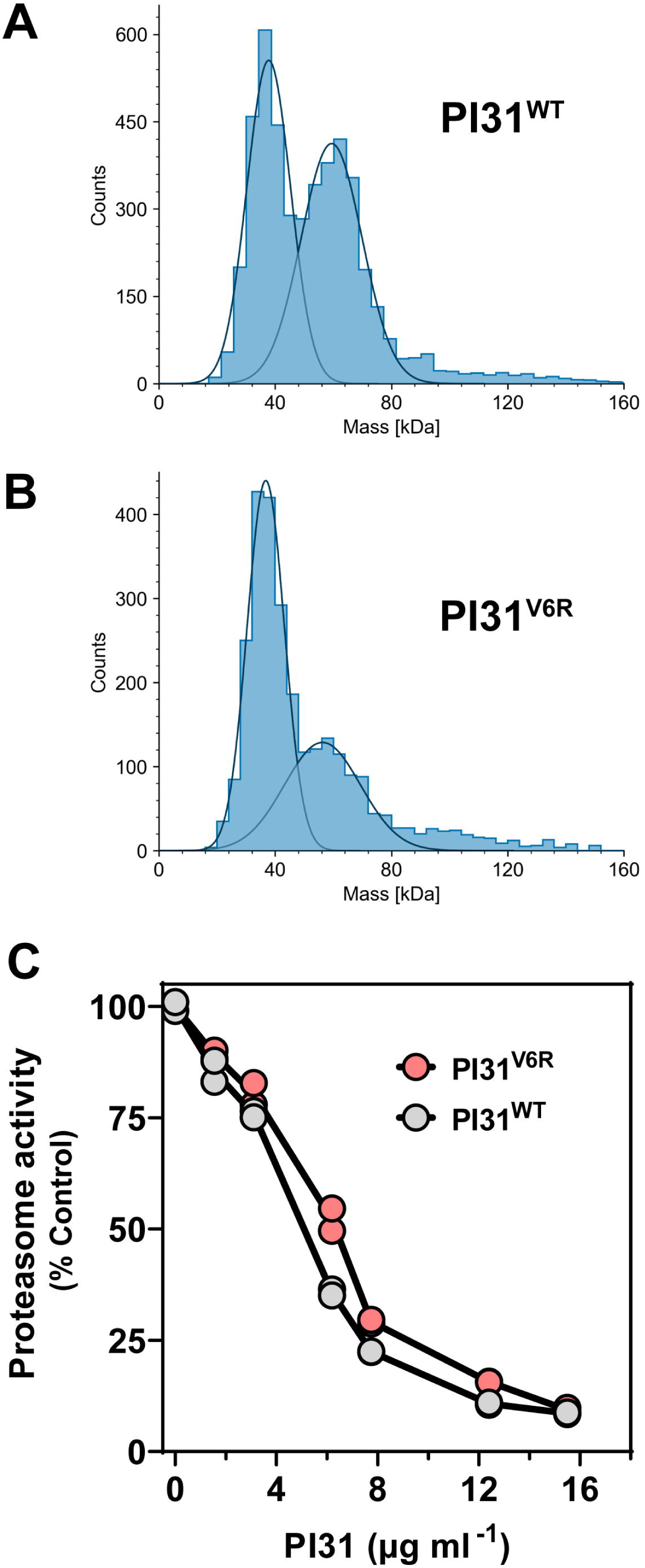
PI31 monomers are sufficient to inhibit 20S proteasome activity. Mass photometry analysis of PI31 dimerization using purified, recombinant (A) hPI31^WT^ and (B) PI31^V6R^ dimerization mutant proteins. Purified recombinant proteins were diluted to 40 nM total monomer (1.2 ng µl^-1^) in PBS during recording and analyzed as described in Experimental Procedures. (C) Inhibition of proteasome activity by PI31^WT^ and dimerization mutant PI31^V6R^. 20S proteasome (50 nM) was activated by 0.03% SDS prior to incubation with a range of PI31 amounts up to 10-fold molar excess. Proteasome activities were measured in duplicate as described in Experimental Procedures, and expressed as percentages relative to controls lacking PI31.

**Figure 8.**
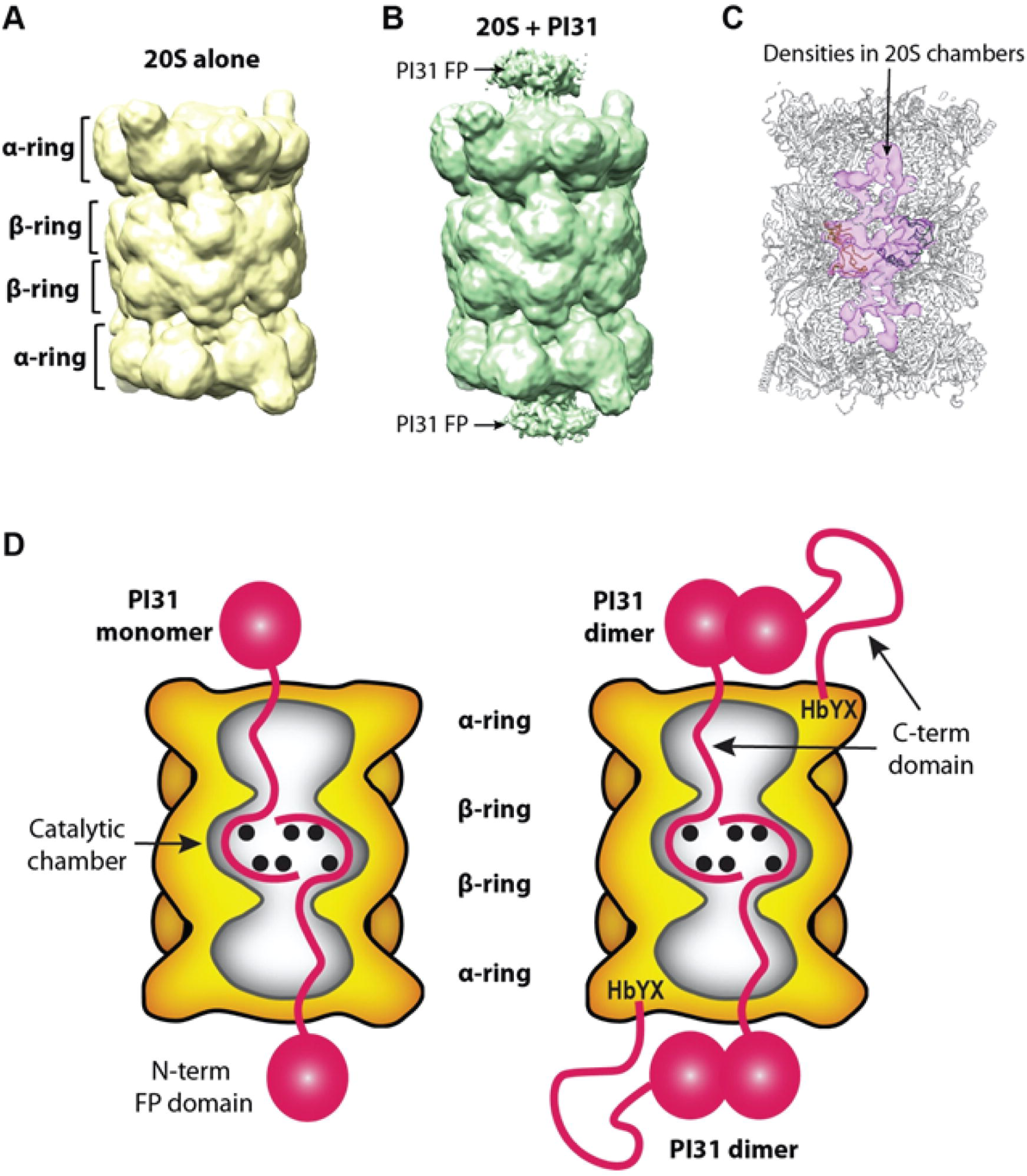
Proposed model for hPI31 interaction with 20S proteasome. A) The low pass filtered EM map of 20S proteasome surface rendered at a low threshold of σ=2. B) The low pass filtered EM map of PI31-bound 20S proteasome surface rendered at a low threshold with σ=2. C) Difference map (pink surface) between the two above low pass filtered EM maps revealing extra PI31 densities inside both catalytic and antechambers. The 20S atomic model is superimposed to indicate the chamber location of the difference densities. D) Alternative models of PI31-20S proteasome complex involving either monomeric (left) or dimeric (right) PI31 showing the interaction of the PI31 C-terminal domain with proteasome catalytic sites (see text for details).

### PI31 overexpression inhibits proteasome activity in mammalian cells

The structure of the PI31-20S complex and consequent proteasome inhibition described above raises questions about their physiologic significance. Previous work by us and others has failed to demonstrate an expected inhibitory effect of PI31 overexpression on proteasome activity in mammalian cells [17, 18]. However, global quantitative proteomic analysis of multiple mammalian cell lines documents that PI31 is normally expressed at low levels in most cells and is exceeded on a molar basis by 20S proteasome by 50-100 fold [33–36]. Retrospective analysis of our previous experiments reveals that PI31 overexpression in those studies was only 3-10 fold over endogenous levels, making it unlikely that any effect on proteasome activity could be detected since PI31 would remain significantly sub-stoichiometric to the proteasome. We have repeated those experiments under conditions that express PI31 by >700 fold over endogenous levels and in >20-fold molar excess of proteasomes (Supplemental Figure 3). Under these conditions, overall proteasome activity in cell extracts was reduced by 25-30% as monitored by proteasome-dependent hydrolysis of peptide substrates (Figure 9). Although additional experiments will be required to determine the exact basis for and specificity of this effect, these data are consistent with a model in which PI31 can bind to and inhibit cellular proteasome.

**Figure 9.**
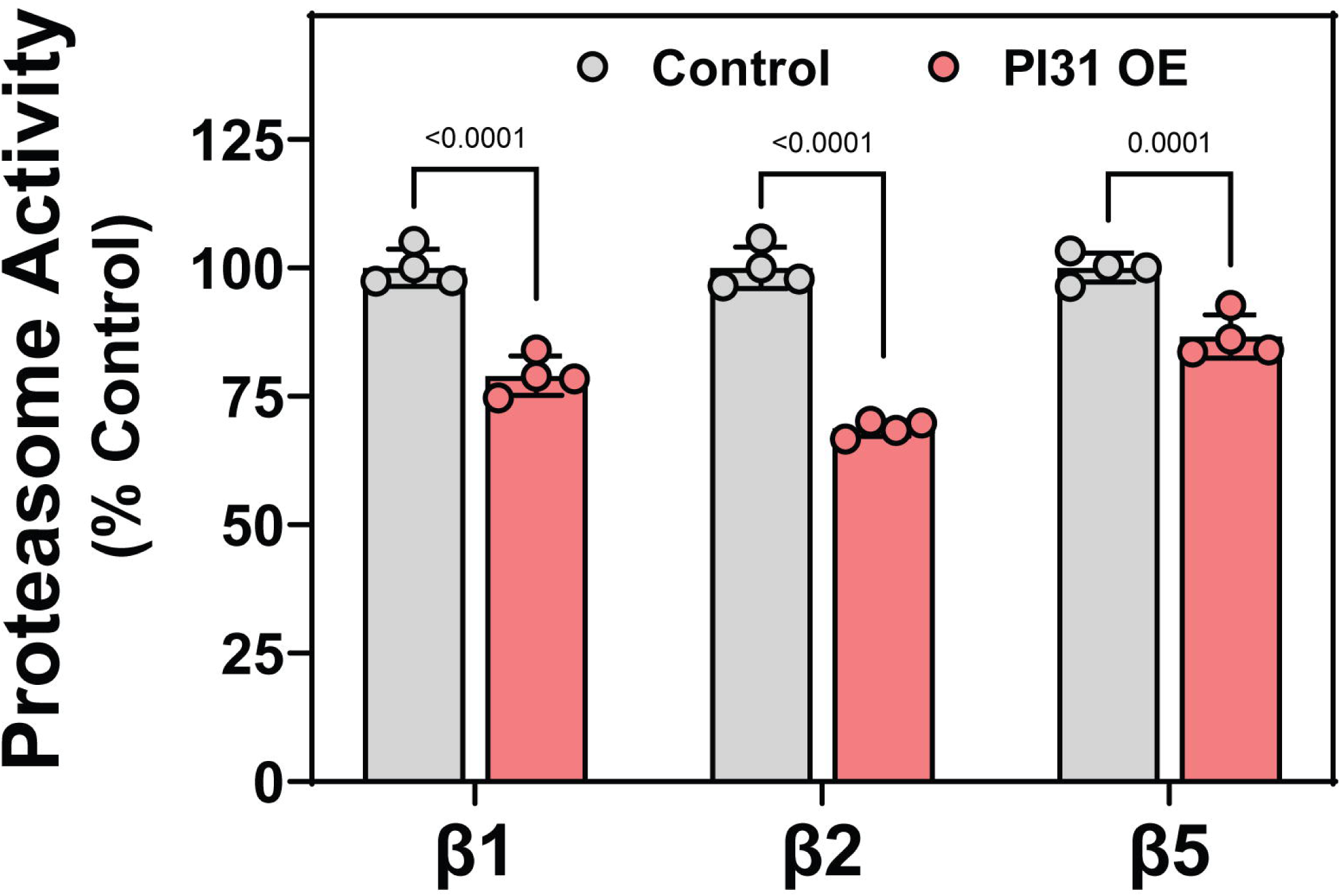
Overexpression of hPI31 inhibits proteasome activity in HepG2 cells. HepG2 cells were transfected with pCMV3-FlagPI31 or vehicle controls as described in Experimental Procedures. Extracts from control and PI31 overexpression conditions were prepared and assayed for proteasome activity using peptide substrates for each of the indicated catalytic subunits, as described in Experimental Procedures. Bars are mean (± SD) activities from n=4 lysates of independently transfected cells. Activity in control (-FlagPI31) extracts was set as 100 and other activities within the same experiment are expressed as a percentage of that value. Differences in activity were analyzed by 2-way ANOVA and multiple comparisons within each catalytic subunit were tested by Sidak’s multiple comparisons test. Similar results were obtained in three independent experiments.

## DISCUSSION

The high-resolution structure of the mammalian PI31-20S proteasome complex reported here provides a structural basis for our original discovery and description of PI31 as an *in vitro* inhibitor of 20S proteasome activity [15]. The structure is in accord with mutational analysis showing that the C-terminal domain of PI31 is necessary and sufficient for inhibitory activity (Figure 5 and [16]). This mammalian structure and two recently published structures of yeast and *Microsporidia* PI31-like proteins in complex with 20S proteasome reveal a surprising mechanism of inhibition via direct interaction of PI31 with the proteasome’s catalytic sites [22]. This mechanism differs significantly from previously anticipated models of inhibition in which the C-terminus of PI31 binds to the exterior of the proteasome near the substrate access pore and sterically blocks the entrance of substrates to the catalytic chamber. Rationale for the latter model was reinforced by recognition that PI31 contains a C-terminal HbYX motif found in various proteasome regulators that bind to HbYX-cognate sites between outer ring α-subunits [17, 37, 38]. However, as discussed below, the functional relationship between the hPI31’s HbYX motif and the structure of the hPI31-20S complex remains uncertain.

Unlike the unstructured proline/glycine-rich C-terminal domain, the N-terminal FP domain of hPI31 is well-folded [32, 39] and therefore cannot transit the narrow substrate access pore regardless of its gating status. Our previous data showed that this domain neither binds to nor inhibits the 20S proteasome [16]. No density for the FP domain was detected in our high-resolution EM map of the hPI31-20S complex, likely due to its flexibility on the exterior of the proteasome. However, a low-pass filter of the map shown at a lower display threshold reveals densities on the apical surface of 20S α-rings of hPI31-bound proteasome (Figure 8A-B). We also examined the difference map between the PI31-bound and PI31-free EM maps and observed hPI31 densities in the 20S antechambers as well as in the catalytic chambers. These findings indicate that the C-terminal regions of PI31 in the catalytic chamber are threaded from the antechambers (Figure 8C). This observation is consistent with the proposed orientation of active site engagement along the primary structure of the hPI31 C-terminus, as described above and discussed further below.

Although the mammalian and yeast PI31/Fub1-20S complexes share general structural features, they have a number of significant distinctions. Importantly, the mammalian complex described here was achieved using native 20S proteasome as opposed to a mutant 20S proteasome with a constitutively open gate. This result demonstrates the capacity of PI31 to inhibit a physiologic form of 20S proteasome normally found in cells and rules out effects that might result from an artificial α-ring. The manner by which the polypeptide chains of hPI31/Fub1 interact with the proteasome’s multiple catalytic sites also differ appreciably. The Fub1 chain traverses a path (in the N-to C direction) from β1 to β5 in the opposing β ring and back to β2 in the original β ring. Moreover, Fub1’s interaction with and proposed inhibition of β5 employs the cooperative action of a second Fub1 molecule. In contrast, our data show a hPI31 interaction path (in the N-to C direction) from β5 to β2 to β1. As with Fub1, hPI31 interacts with β5 on the opposing ring from that of its interactions with β1 and β2. However, unlike Fub1, hPI31 utilizes two different segments of the same polypeptide chain for its β5 interaction. This feature indicates that a single molecule of hPI31 is sufficient to engage one copy of each β-type catalytic subunit. The functional consequence of inhibition of only one of two copies of each catalytic subunit type is unknown. Interestingly, an interaction path similar to that of the mammalian proteins has been observed for a PI31-like peptide and 20S proteasome from *Microsporidia* [23]. Nevertheless, the finding that PI31 orthologs from evolutionally diverse organisms directly contact proteasome catalytic sites suggests a highly conserved *in vivo* inhibitory function for this interaction.

hPI31 can form homodimers mediated by the N-terminal FP domain [16, 32, 39] but exists in a monomer-dimer equilibrium at PI31 concentrations likely found in cells (Figure 7 and [16]). We favor a model in which the hPI31-20S complex results from 20S interaction with PI31 monomers (Figure 8D). This feature might contribute to the observation that high molar ratios of PI31 to 20S are required for inhibition *in vitro*. It is also consistent with the conclusion that the two C-termini residing the catalytic chamber enter from opposite ends of the proteasome. Nevertheless, we cannot exclude models of inhibition involving PI31 dimers (Figure 8D). In one variant of this model, transit through the substrate access pore may be limited to one of the two C-termini while the HbYX motif of the other subunit promotes gate opening via α-subunit binding. Although such a mechanism is consistent with previously shown gate-opening properties of hPI31 C-terminal peptides, it cannot be obligatory because PI31 lacking HbYX residues retains inhibitory activity (Figure 5 and [16, 17]). Alternatively, both C-termini of a PI31 dimer may enter the antechamber through the same pore while only one gains access to the central catalytic chamber. Interestingly, previous work has shown that the open proteasome pore can accommodate two unstructured polypeptide chains [40]. Additional work will be required to distinguish among these and related models and to determine their functional implications.

In its position on the apical surface of the α-rings, the FP domain of PI31 may serve as a binding partner of other proteins. In fact, FbxO7, an F-box protein whose mutant forms are responsible for early-onset autosomal recessive Parkinson’s disease, has been identified as a PI31 binding partner and this interaction is mediated by the FP domains of both proteins [32, 41, 42]. Likewise, the FP domain is likely involved in a proposed role for PI31 as an adaptor between the proteasome and motor proteins for axonal transport of proteasomes to intracellular sites of action [21]. The relationship between PI31-mediated proteasome inhibition and such functions is unclear and would appear to require release of inhibition upon delivery. Although we have not identified *in vitro* conditions that reverse PI31-proteasome binding and inhibition, such a process has been proposed as a mechanism for termination of PI31-like mediated proteasome inhibition that allows assembly of active 20S-19S/PA700 holoenzyme complexes after germination of dormant *Microporidia* spores [23]. The generality of the latter process is unknown and appears to be inconsistent with previous evidence that PI31 blocks the *in vitro* assembly and activation of proteasome holoenzymes with 19S/PA700 and PA28 regulators [16, 43]. Alternatively, irreversible PI31 binding could result in clearance of aberrant or unneeded proteasomes, as was hypothesized for Fub1. Such a process might require binding of the FP domain with proteins involved in proteasome disposal and is consistent with a general role for the exposed FP domain in protein-protein interactions.

Because hPI31 normally is present in cells at much lower levels than the proteasome, it can affect only a small proportion of total cellular proteasome. Therefore, any physiologic effect of even full-engaged PI31 would escape detection in assays that measure global proteasome activity. In contrast, we have shown that proteasome activity was detectably inhibited when PI31 was ectopically expressed to levels exceeding those of the proteasome (Figure 9). Although additional work is required to determine the exact mechanism of this effect, it is consistent with a direct inhibitory role for cellular PI31. We are unaware of any physiologic or pathologic condition in which PI31 levels are increased to such levels. Nevertheless, endogenous levels of PI31 may serve to terminate the activity of a subpopulation of proteasomes that function normally in a regulator-free fashion [44–47], or to prevent dysregulation of cellular proteostasis by inappropriately activated 20S proteasomes [22, 45].

## EXPERIMENTAL PROCEDURES

### Protein purification

20S proteasome was purified from bovine red blood cells as described previously [29]. Recombinant wild-type human PI31 and PI31 truncation mutants were expressed in *E.coli* and purified as described previously [16]. A pET28a(+) vector containing N-terminally tagged, wild type hPI31 was utilized for site directed mutagenesis. Primers and annealing temperatures for site-directed mutagenesis were predicted using the NEBaseChanger online tool (version 1.3.3, https://nebasechanger.neb.com/) . Site-directed mutagenesis was performed on 10 ng pET28a(+)-HisPI31^wt^ template with the Q5 Site-Directed Mutagenesis Kit (New England BioLabs) according to manufacturer’s instructions. Based on the length of the plasmid, an extension time of 2.5 minutes was used for each PCR cycle. The mutagenesis PCR product was ligated using the Kinase Ligase DpnI (KLD) mix provided in the kit. Ligated plasmid (1 µl) was then transformed into One Shot BL21 Star (DE3) competent cells (Invitrogen) with outgrowth in SOC media for 1 hour at 37°C before plating on LB agar plates containing kanamycin (50 µg/ml). Sequences were confirmed by Sanger sequencing at the UT Southwestern McDermott Center Sequencing Core with Big Dye Terminator 3.1 and capillary instruments (Applied Biosystems Inc.).

Recombinant His-PI31 mutant proteins were purified by metal affinity chromatography. BL21 Star (DE3) cells containing pET28a(+) HisPI31 mutants from monoclonal stab cultures were used to inoculate 150 ml LB broth containing 50 µg/ml kanamycin. Cells were grown with shaking overnight at 37°C. The following morning, 100 ml of LB/Kan broth was added to cultures with 250 – 500 µM IPTG to induce expression. Cells were grown at 37°C for an additional 3-6 hours. Following expression, cells were pelleted by centrifugation at 10,000 x g for 10 minutes at 4°C. Cell pellets were resuspended in 8 ml Binding Buffer (50 mM sodium phosphate, pH 7.6, 500 mM NaCl, 10 mM imidazole, pH 8.0) containing 0.1 mg/ml lysozyme and 1x cOmplete, EDTA-free protease inhibitor cocktail (Roche). Cells were incubated on ice for 30 minutes before sonication. Lysed cells were then centrifuged at 14,000 rpm, 20 minutes, 4°C. The 8 ml supernatant was applied to 1 ml packed volume of Ni-NTA His-Bind Resin (Millipore Sigma). Binding was performed with gentle mixing for 1 hour at 4°C and loaded onto a Poly Prep Chromatography Column (BioRad). The loaded resin was then washed 3-4 times with 6 ml Wash Buffer (50 mM sodium phosphate, pH 7.6, 500 mM NaCl, 20 mM imidazole, pH 8.0) until protein was undetectable with Bradford Reagent (Pierce). His-PI31 was eluted in four 0.5 ml fractions with 250 mM imidazole, 50 mM sodium phosphate, pH 7.6, 500 mM NaCl. Fractions containing peak protein (as assessed by Bradford assay) were pooled and dialyzed against Buffer H (20 mM Tris-HCl, pH 7.6 at 4°C, 20 mM NaCl, 1 mM EDTA, 5 mM βME) containing 10% glycerol. Purification was confirmed by SDS-PAGE Coomassie staining and Western blotting with anti-His antibody.

### Proteasome activity assays

Inhibitory activity of hPI31 proteins against bovine and human 20S proteasome was analyzed using 7-amino-4-methylcoumarin (amc)-linked fluorogenic peptide substrates specific to each of the proteasome catalytic sites: caspase-like (β1, Z-LLE-amc); trypsin-like (β2, Ac-RLR-amc or Z-VVR-amc); and chymotrypsin-like (β5, Suc-LLVY-amc). In some experiments, latent 20S proteasome was first activated with a pre-incubation in 0.03% SDS at 37°C for 5 minutes. PI31 (wild-type, D197A, or D244A) was then for an additional 15 minutes at 37°C. Proteasome activity was assayed by addition of peptide substrate (50 µM final concentration). Free amc produced by proteasome hydrolysis was monitored (Ex. 360 nm, Em. 460 nm) at 45 second intervals over 21 minutes at 37°C (Synergy Mx Multi-Mode Microplate reader, BioTek). Proteasome activity was expressed as rates of amc production and all were linear with respect to time and proteasome concentration in these assays. All assays were performed in triplicate and repeated independently at least three times. Differences between the effects of PI31 mutants on proteasome activity were analyzed by one-way ANOVA and Tukey’s HSD test using GraphPad Prism 9 (v9.5.0). PI31 mutant inhibitory activity was also assessed using direct fluorescent labeling of proteasome active sites with a Me_4_BodipyFL-Ahx_3_Leu_3_-VS activity-based probe [30]. This probe covalently binds each of the β1, β2, and β5 active sites and can be visualized following separation by SDS-PAGE. Proteasomes were first SDS-activated and incubated with PI31 as described above. Activity-based probe (0.5 - 1 µM final concentration) was then added for an additional 15-40 min incubation at 37°C. Because the probe labels each subunit with different affinity, concentrations and incubation times were adjusted to avoid signal saturation of some subunits while maintaining detectability of others.

Labelling was quenched by denaturing proteasomes in SDS-PAGE sample loading dye and proteins were separated on a SDS-PAGE gel. Gels were either imaged directly using an Odyssey M Imager (LI-COR, 520 nm fluorescence channel) or transferred to nitrocellulose membranes and visualized on ChemiDoc MP Imager (BioRad, fluorescein channel). Images were imported into Image Studio (LI-COR, v5.0.21) for quantification.

### Western blotting and antibodies

Western blotting was conducted as described previously [17]. Transferred membranes were by imaged using a LiCor Odyssey CLx. Rabbit polyclonal antibody against human PI31 was purchased from Enzo (PW9710) and was used at 1:2000 dilution. Rabbit polyclonal antibody against human the β5 subunit of the proteasome was produced and characterized in the DeMartino lab as described previously [48, 49]. Rabbit polyclonal antibody against human β tubulin was purchased from Cell Signaling Technology (2146) and used at 1:1000 dilution. Goat anti-rabbit IRDye680RD was purchased from Li-COR and was used at 1:10000 dilution.

### Overexpression of PI31 in mammalian cells

A mammalian expression vector, pCMV3-N- FlagPI31^wt^ was obtained from Sino Biological Inc (catalog #HG17079-NF). HepG2 cells were seeded in 6-well plates at 3 × 10^5^ cells well^-1^ and grown overnight at 37°C, 5% CO_2_ in DMEM media containing 10% FBS, penicillin, and streptomycin. The following day, 750 ng of pCMV3- N-FlagPI31 vector was mixed with 5 µl P3000 reagent diluted in OptiMEM media and added to 5 µl of diluted Lipofectamine3000 according to manufacturer’s instructions (Life Technologies). Vectors or vehicle-only controls were added to replicate wells and transfected for 24 hours. Following transfection, cells were washed three times with PBS, and harvested in a hypotonic lysis buffer in the presence of ATP (20 mM Tris, pH 7.6 at 37°C, 20 mM NaCl, 5 mM MgCl_2_, 100 µM ATP, 1 mM DTT, 0.1% NP-40). Cells were then passed through a 27 ½ G needle 15 times and centrifuged at 14,000 rpm, 20 min, 4°C. Supernatants were divided into three sets of triplicate wells in a 96-well plate and assayed for proteasome activity using fluorogenic peptide substrates (β1, Z-LLE-amc; β2, Z-VVR-amc; and β5, Suc-LLVY-amc) as described above for pure proteins.

Proteasome and PI31 amounts in HepG2 cells transfected with vehicle controls or pCMV3- N-FlagPI31 were analyzed by semi-quantitative Western blotting using polyclonal antibodies against the proteasome β5 subunit and the PI31 C-terminus. Within the same gel, a range of pure proteins (bovine 20S proteasome or recombinant His-PI31) was used as standards for quantification. Secondary antibodies with IRDye (LI-COR) were used and NIR signals were quantified on Odyssey CLx imager (LI-COR). Signals were converted to protein masses based on the standard curve of pure proteins, and then converted to moles assuming values of 700 kDa for proteasome and 31 kDa for PI31 (Supplemental Figure 3). Amounts of each protein were then normalized to total protein content (determined by BCA) to compare molar ratios of PI31 and proteasome in HepG2 cells.

### Analysis of PI31 oligomerization by mass photometry

Purified His-PI31 (wild-type or V6R mutant) proteins were dialyzed against 20 mM Tris HCl (pH 7.6 at 5°C), 20 mM NaCl for analysis of oligomerization by mass photometry (TwoMP, Refeyn). Protein concentrations were determined by BCA assay (Pierce), and converted to molar concentrations assuming 31 kDa molecular weights. PI31 was then diluted in 0.2 µm filtered PBS into a total monomer concentration of 400 nM. Cover slips were cleaned with isopropanol and MilliQ water and fitted with silicon gaskets. The mass photometer optics were focused on 16.2 µl buffer prior to the addition and mixture of 1.8 µl sample. Sample mixture into the pre-focused droplet resulted in an additional 1/10 dilution (40 nM final), and recordings were initiated immediately after mixing. Bovine serum albumin (BSA) and thyroglobulin oligomers were used as a molecular weight standard to calibrate ratiometric contrast measurements to molecular weight. Automatic Gaussian curve fitting for peaks was performed by DiscoverMP software (Refeyn) which identified peaks centered at 36-37 and 56-59 kDa for monomers and dimers, respectively.

### Cryo-EM sample preparation and data collection

Cryo-EM was conducted at the Cryo-EM facility of Van Andel Institute. To obtain the human PI31-bovine proteasome complex, we mixed purified hPI31 and 20S at a molar ratio of 5:1 and incubated the mixture at 37°C for 1 hour in 20 mM Tris, pH 7.5, 5 mM MgCl_2_, 100 mM KCl, and 1mM DTT. The sample concentration was adjusted to 3 mg/mL for EM grid preparation. Cryo-EM grids were prepared in Vitrobot Mark IV (FEI, USA) with the chamber set to 5°C and 100% humidity. We first glow-discharged Quantifoil R 1.2/1.3 300 mesh gold holey carbon grids with O_2_-air for 30 s at 30 W in a Gatan Model 950 Advanced Plasma System, applied a droplet of 3 µl sample to the EM grid, waited for 1s, blotted the grid for 4 s using a piece of 595 filter paper with a blotting force set to 3, then vitrified the sample by plunging the grid into liquid ethane precooled with liquid nitrogen. Cryo-EM dataset was recorded in a Titan Krios (FEI, USA) equipped with a Quantum 967 energy Filter plus a post GIF K3 summit direct electron detector (A.K.A. BioQuantum). Movies were recorded using SerialEM software in super-resolution counting mode with a defocus range of -1.3 to -1.8 μm and a pixel size of 0.414 Å/pixel. The dose rate was 46 electrons/Å^2^/s with 1.3 s exposure time for a total dose of 60 electrons/Å^2^. A total of 19,264 movies were collected during a two-day session.

### Cryo-EM data processing

Image processing was performed in Relion 3.1.1 and Cryosparc 3.3.2 [50, 51]. We first corrected the beam-induced motion of each movie micrograph using MotionCor2 with binning factor of 2 in Relion [52]. The corrected micrographs were imported into Cryosparc for further processing. We used CTFFIND4 to estimate the defocus value for each micrograph and used the detected value to correct for the contrast transfer effect. Based on the CTF detection limit, low resolution micrographs were removed, and a total of 18,528 micrographs were retained for particle picking. We first used blob picking to generate several templates for the next stage template-based particle picking. We used the proteasome particles from template picking and with good 2D class averages to train Topaz and extracted 1,573,299 particles by the trained Topaz. After reference-free 2D classification and subsequent 3D *ab initio* 3D reconstruction, a total of 1,467,640 particles were selected for 3D multi-reference heterogenous refinement. The 3D class with extra density inside the catalytic β-ring of the 20S proteasome was chosen for further 3D heterogenous refinement using the classified maps with and without extra density as the references. The particles belonging to the two 3D classes (with or without extra density) were segregated and refined separately, first by homogeneous refinement and followed by non-uniform refinement [50]. By applying C1 symmetry during refinement, we obtained a 3D EM map of the 20S proteasome with extra chamber density (PI31) at 2.54 Å average resolution. We applied C2 symmetry to the 20S proteasome alone particles and refined a 3D EM map at 2.23 Å average resolution. The resolutions were estimated by applying a soft mask around the map of proteasome and used the gold standard Fourier shell correlation (FSC) =0.143 as the threshold.

### Model building

Both 20S alone and PI31-20S complex were modeled using the coordinates of bovine 20S proteasome (PDB ID 1IRU) as the initial model. The PI31 structure was built manually in Coot after completion of the 20S proteasome modeling [53]. Real space refinements were performed in Phenix to correct geometric errors, and the model quality was estimated using MolProbity in Phenix [54].

## SUPPORTING INFORMATION

This article contains supporting information (Supplemental Figures 1-3 and Supplemental Table 1)

## Acknowledgements

Cryo-EM dataset were collected at the David Van Andel Advanced Cryo- Electron Microscopy Suite in Van Andel Institute. We thank G. Zhao and X. Meng for facilitating data collection. This work was supported by the US National Institutes of Health grants AI070285 (to H.L.) and GM129088 (to G.N.D.) and the Van Andel Institute (to H.L.). PI31 oligomerization data were obtained using a mass photometer that was supported by award S10OD030312-01 from the National Institutes of Health.

## Author contributions

H.L. and G.N.D. designed research; H.H. performed cryo-EM, image processing, 3D reconstruction, and atomic modeling. J.W. and A.K. performed biochemical and cellular analyses. H.H., J.W., H.L, and G.N.D. wrote the manuscript with input from all authors.

## Competing interests

The authors declare no competing interests

**Supplemental Figure 1. Cryo-EM of the hPI31-20S complex.** A) A representative micrograph after motion correction. B) Selected 2D class averages. C-D) Local resolution estimation of the final maps of the bovine 20S in the absence of (C) and presence of PI31 (D). E-F) Gold standard Fourier shell correlation curve for the maps of PI31-free 20S (E) and the PI31-bound 20S (F). The resolution was determined at FSC=0.143.

**Supplemental Figure 2. The flow chart of cryo-EM data processing.** Starting from a single dataset collected from the mixture of the PI31 and 20S, we segregated the particle images into two major classes – 20S CP with and without the bound PI31, leading to the two final EM maps of 20S CP particle alone at 2.23 Å average resolution and the 20S complexes with PI31 at 2.54 Å average resolution.

**Supplemental Figure 3. Quantification of relative contents of PI31 and 20S proteasome in HepG2 cells with or without overexpression of hPI31.** HepG2 cells were transfected with pCMV3-FlagPI31 or vehicle controls as described in Experimental Procedures. Cell extracts matching those for which data are reported in Figure 9 were blotted for 20S proteasome (A) or PI31 (B). Relevant proteins were quantified for each cellular condition using pure proteins as standards within the same blot. C) Relative levels of cellular 20S proteasome and PI31 under control and overexpression condition in four independent samples.

## Notes

### Competing Interest Statement

The authors have declared no competing interest.

### Summary of Updates

Correct order of authors. Replace Figure 8 with corrected figure.

## REFERENCES

1. Hanna, J., et al., Protein Degradation and the Pathologic Basis of Disease. Am J Pathol, 2019. 189(1): p. 94–103.

2. Lecker, S.H., A.L. Goldberg, and W.E. Mitch, Protein degradation by the ubiquitin-proteasome pathway in normal and disease states. J Am Soc Nephrol, 2006. 17: p. 1807–1819.

3. Tanaka, K. and N. Matsuda, Proteostasis and neurodegeneration: the roles of proteasomal degradation and autophagy. Biochim. Biophys. Acta, 2014. 1843(1): p. 197–204.

4. Türker, F., E.K. Cook, and S.S. Margolis, The Proteasome and its Role in the Nervous System. Cell Chem Biol, 2021. 28(7): p. 903–17.

5. Rock, K.L., et al., Inhibitors of the proteasome block the degradation of most cell proteins and the generation of peptides presented on MHC class I molecules. Cell, 1994. 78: p. 761–771.

6. DeMartino, G.N. and T.G. Gillette, Proteasomes: machines for all reasons. Cell, 2007. 129: p. 659–662.

7. Groll, M., et al., Structure of the 20S proteasome from yeast at 2.4A resolution. Nature, 1997. 386: p. 463–471.

8. Baumeister, W., et al., The proteasome: paradigm of a self-compartmentalizing protease. Cell, 1998. 92: p. 367–380.

9. Bochtler, M., et al., The proteasome. Ann. Rev. Biophys. Biomol. Struct, 1999. 28: p. 295–317.

10. Collins, G.A. and A.L. Goldberg, The Logic of the 26S Proteasome. Cell, 2017. 169(5): p. 792–806.

11. Finley, D., X. Chen, and K.J. Walters, Gates, Channels, and Switches: Elements of the Proteasome Machine. Trends Biochem Sci, 2016. 41(1): p. 77–93.

12. Whitby, F.G., et al., Structural basis for the activation of 20S proteasomes by 11S regulators. Nature, 2001. 408: p. 115–120.

13. Smith, D.M., et al., Docking of the proteasomal ATPases’ carboxyl termini in the 20S proteasome’s alpha ring opens the gate for substrate entry. Mol. Cell, 2007. 27(5): p. 731–744.

14. Sakata, E., M.R. Eisele, and W. Baumeister, Molecular and cellular dynamics of the 26S proteasome. Biochim Biophys Acta Proteins Proteom, 2021. 1869(3): p. 140583.

15. Ma, C.-P., C.A. Slaughter, and G.N. DeMartino, Purification and characterization of a protein inhibitor of the 20S proteasome (macropain). Biochim. Biophys. Acta, 1992. 1119: p. 303–311.

16. McCutchen-Maloney, S.L., et al., cDNA cloning, expression, and functional characterization of PI31, a proline-rich inhibitor of the proteasome. J. Biol. Chem, 2000. 275: p. 18557–18565.

17. Li, X., et al., Molecular and cellular roles of PI31 (PSMF1) protein in regulation of proteasome function. J. Biol. Chem, 2014. 289(25): p. 17392–17405.

18. Zaiss, D.M., et al., PI31 is a modulator of proteasome formation and antigen processing. Proc. Natl. Acad. Sci. U. S. A, 2002. 99(22): p. 14344–14349.

19. Bader, M., et al., A conserved F-box–regulatory complex controls proteasome activity in Drosophila. Cell, 2011. 145(3): p. 371–82.

20. Cho-Park, P.F. and H. Steller, Proteasome Regulation by ADP-Ribosylation. Cell, 2013. 153(3): p. 614–627.

21. Liu, K., et al., PI31 Is an Adaptor Protein for Proteasome Transport in Axons and Required for Synaptic Development. Dev Cell, 2019. 50(4): p. 509–524.e10.

22. Rawson, S., et al., Yeast PI31 inhibits the proteasome by a direct multisite mechanism. Nat Struct Mol Biol, 2022. 29(8): p. 791–800.

23. Jespersen, N., et al., Structure of the reduced microsporidian proteasome bound by PI31-like peptides in dormant spores. Nat Commun, 2022. 13.

24. Löwe, J., et al., Crystal structure of the 20S proteasome from the archaeon T. acidophilum at 3.4 Ä resolution. Science, 1995. 268: p. 533–539.

25. Groll, M., et al., The catalytic sites of 20S proteasomes and their role in subunit maturation: A mutational and crystallographic study. Proc Natl Acad Sci U S A, 1999. 96(20): p. 10976–83.

26. Keil, B., Essential substrate residues for action of endopeptidases, in Specificity of proteolysis. 1992, Springer. p. 43–228.

27. Religa, T.L., R. Sprangers, and L.E. Kay, Dynamic regulation of archaeal proteasome gate opening as studied by TROSY NMR. Science, 2010. 328(5974): p. 98–102.

28. Osmulski, P.A., M. Hochstrasser, and M. Gaczynska, A tetrahedral transition state at the active sites of the 20S proteasome is coupled to opening of the alpha-ring channel. Structure, 2009. 17(8): p. 1137–1147.

29. McGuire, M.J., et al., The high molecular weight multicatalytic proteinase, macropain, exists in a latent form in human erythrocytes. Biochim. Biophys. Acta, 1989. 995: p. 181–186.

30. Schipper-Krom, S., et al., Visualizing Proteasome Activity and Intracellular Localization Using Fluorescent Proteins and Activity-Based Probes. Front Mol Biosci, 2019. 6.

31. Kisselev, A.F., et al., Proteasome active sites allosterically regulate each other, suggesting a cyclical bite-chew mechanism for protein breakdown. Mol Cell, 1999. 4(3): p. 395–402.

32. Kirk, R., et al., Structure of a conserved dimerization domain within the F-box protein Fbxo7 and the PI31 proteasome inhibitor. J. Biol. Chem, 2008. 283(32): p. 22325–22335.

33. Schwanhäusser, B., et al., Global quantification of mammalian gene expression control Corrigendum: Global quantification of mammalian gene expression control. Nature, 2011. 473(7347): p. 337–42.

34. Beck, M., et al., The quantitative proteome of a human cell line. Mol Syst Biol, 2011. 7: p. 549.

35. Itzhak, D.N., et al., Global, quantitative and dynamic mapping of protein subcellular localization. Elife, 2016. 5.

36. Kulak, N.A., et al., Minimal, encapsulated proteomic-sample processing applied to copy-number estimation in eukaryotic cells. Nature Methods, 2014. 11(3): p. 319–324.

37. Kim, Y.C. and G.N. DeMartino, C termini of proteasomal ATPases play nonequivalent roles in cellular assembly of mammalian 26 S proteasome. J. Biol. Chem, 2011. 286(30): p. 26652–26666.

38. Rabl, J., et al., Mechanism of gate opening in the 20S proteasome by the proteasomal ATPases. Mol Cell, 2008. 30(3): p. 360–8.

39. Shang, J., X. Huang, and Z. Du, The FP domains of PI31 and Fbxo7 have the same protein fold but very different modes of protein-protein interaction. J Biomol Struct Dyn, 2015. 33(7): p. 1528–38.

40. Liu, C.-W., et al., Endoproteolytic activity of the proteasome. Science, 2003. 299: p. 408–411.

41. Di Fonzo, A., et al., FBXO7 mutations cause autosomal recessive, early-onset parkinsonian-pyramidal syndrome. Neurology, 2009. 72(3): p. 240–5.

42. Vingill, S., et al., Loss of FBXO7 (PARK15) results in reduced proteasome activity and models a parkinsonism-like phenotype in mice. Embo j, 2016. 35(18): p. 2008–25.

43. Zaiss, D.M., et al., The proteasome inhibitor PI31 competes with PA28 for binding to 20S proteasomes. FEBS Lett, 1999. 457(3): p. 333–338.

44. Asher, G., N. Reuven, and Y. Shaul, 20S proteasomes and protein degradation “by default”. BioEssays, 2006. 28(8): p. 844–849.

45. Baugh, J.M., E.G. Viktorova, and E.V. Pilipenko, Proteasomes can degrade a significant proportion of cellular proteins independent of ubiquitination. J. Mol. Biol, 2009. 386(3): p. 814–827.

46. Kumar Deshmukh, F., et al., The Contribution of the 20S Proteasome to Proteostasis. Biomolecules, 2019. 9(5).

47. Ramachandran, K.V. and S.S. Margolis, A mammalian nervous system-specific plasma membrane proteasome complex that modulates neuronal function. Nat Struct Mol Biol, 2017. 24(4): p. 419–30.

48. Fabunmi, R.P., et al., Interferon γ regulates accumulation of the proteasome activator PA28 and immunoproteasomes at nuclear PML bodies. J. Cell Sci, 2001. 114: p. 29–36.

49. Kim, Y.C., et al., ATP binding by proteasomal ATPases regulates cellular assembly and substrate-induced functions of the 26 S proteasome. J. Biol. Chem, 2013. 288(5): p. 3334–3345.

50. Punjani, A., et al., cryoSPARC: algorithms for rapid unsupervised cryo-EM structure determination. Nat Methods, 2017. 14(3): p. 290–296.

51. Zivanov, J., et al., New tools for automated high-resolution cryo-EM structure determination in RELION-3. Elife, 2018. 7.

52. Zheng, S.Q., et al., MotionCor2: anisotropic correction of beam-induced motion for improved cryo-electron microscopy. Nat Methods, 2017. 14(4): p. 331–332.

53. Emsley, P., et al., Features and development of Coot. Acta Crystallogr D Biol Crystallogr, 2010. 66(Pt 4): p. 486–501.

54. Liebschner, D., et al., Macromolecular structure determination using X-rays, neutrons and electrons: recent developments in Phenix. Acta Crystallogr D Struct Biol, 2019. 75(Pt 10): p. 861–877.

