## Supplementary figures and images for "High-resolution structure of mammalian PI31–20S proteasome complex reveals mechanism of proteasome inhibition"

### Supplemental Figure 1

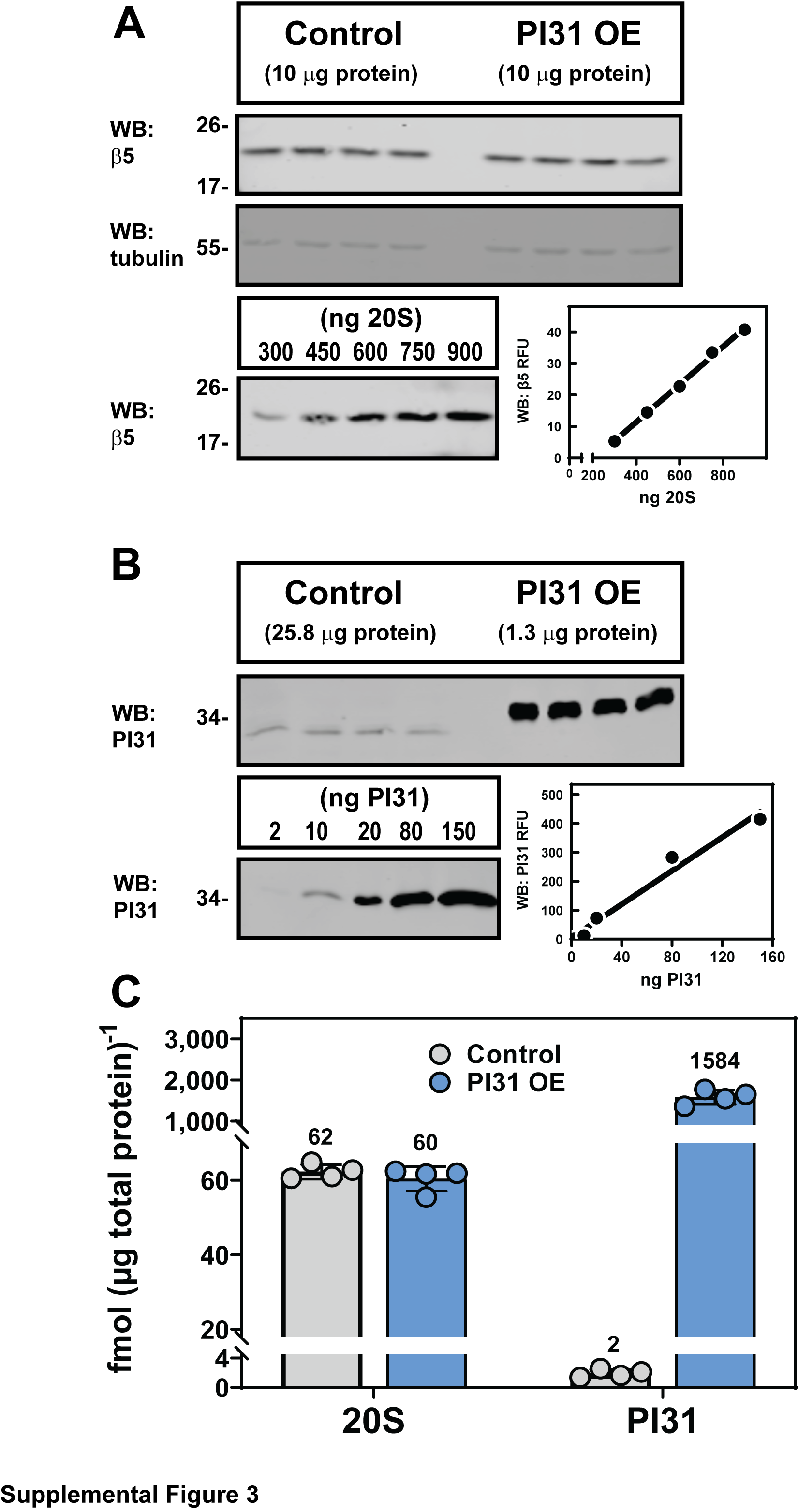

### Supplemental Figure 2

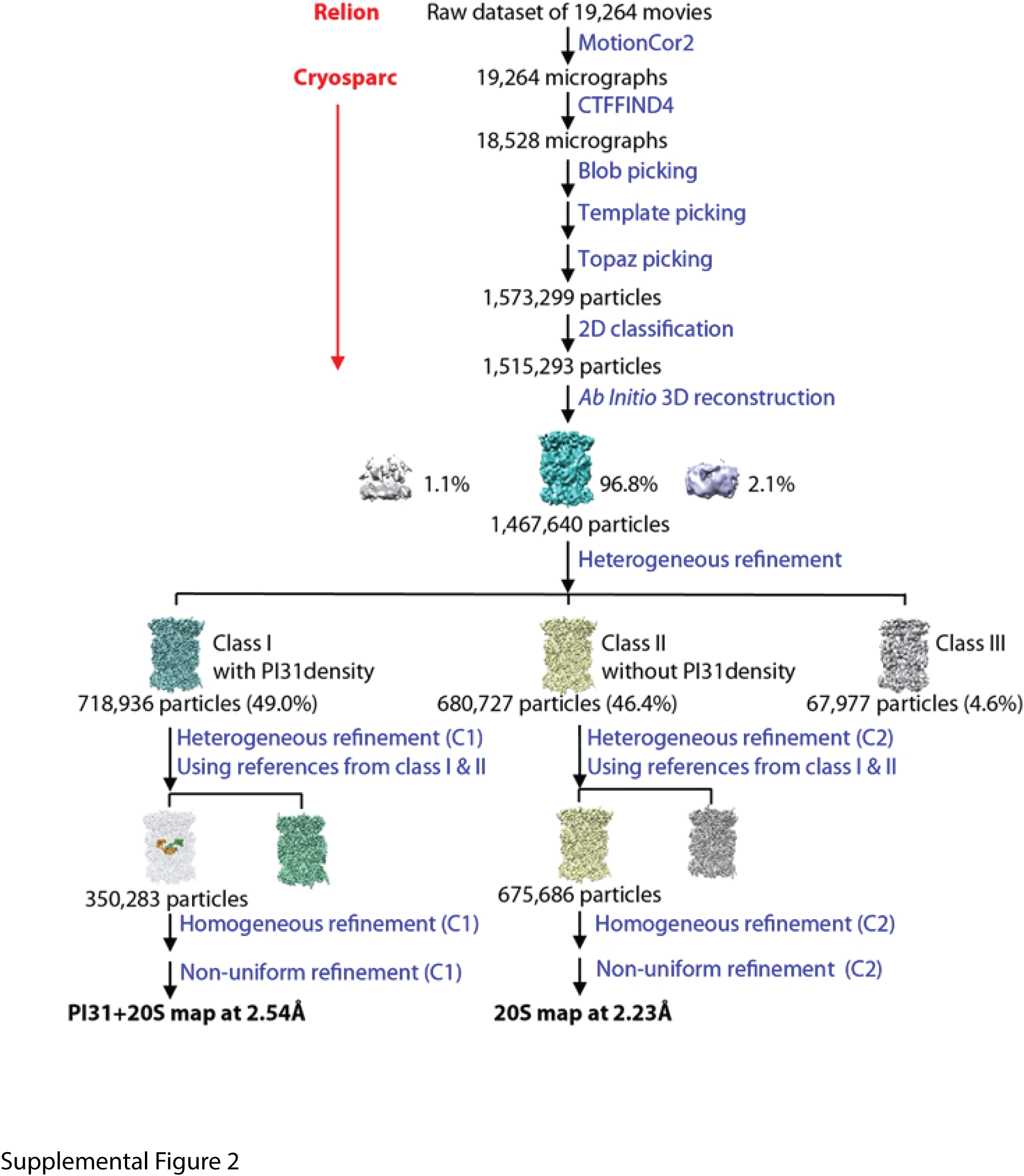

### Supplemental Figure 3

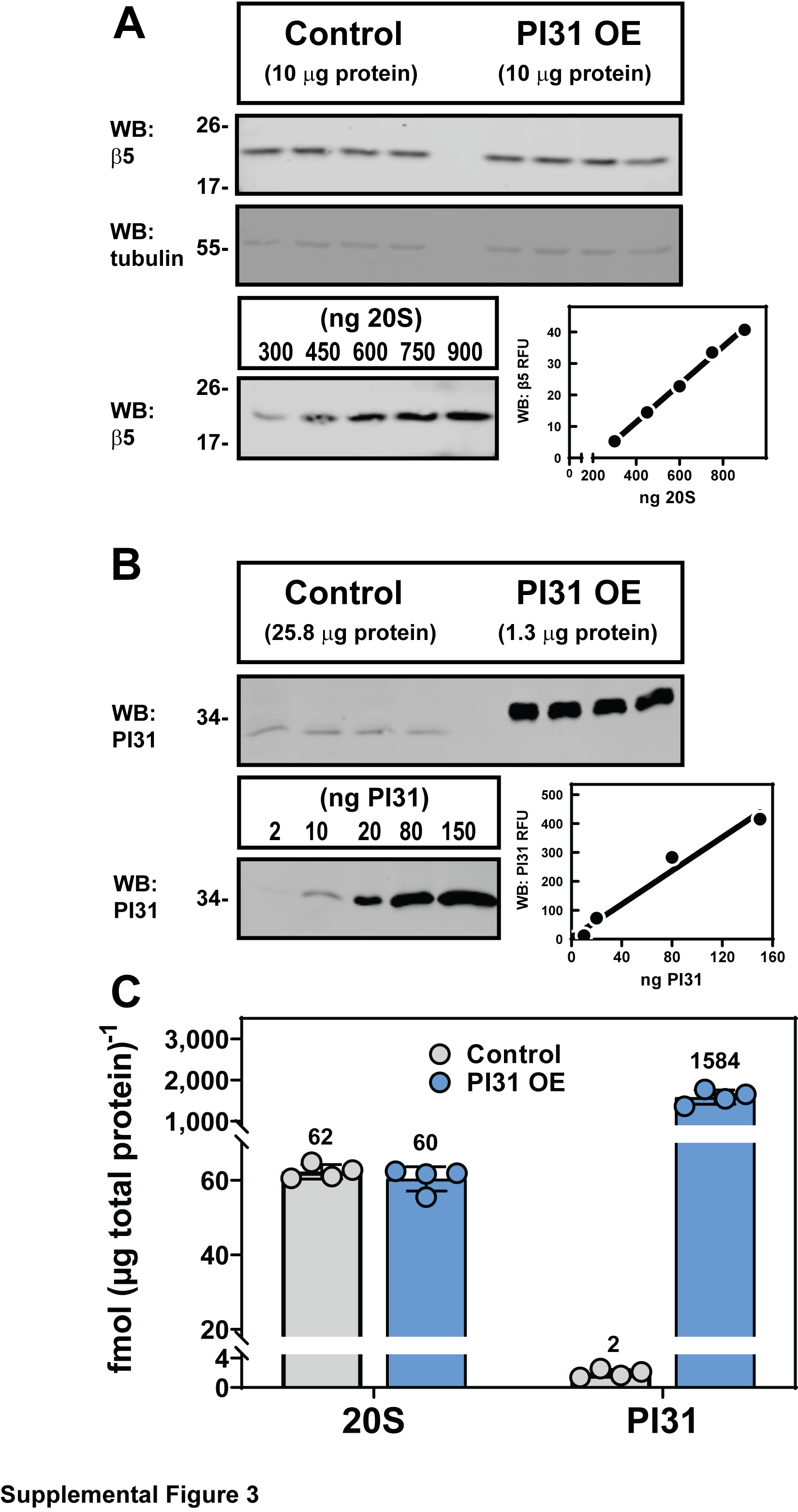
