## Supplemental Table 1 for "High-resolution structure of mammalian PI31–20S proteasome complex reveals mechanism of proteasome inhibition"

**Supplemental Table 1. Cryo-EM data collection, refinement, and atomic model validation**

|  | Bovine 20S proteasome | Bovine 20S complexed  with human PI31 |
| --- | --- | --- |
| PDB code | 8FZ5 | 8FZ6 |
| Data collection and processing |  |  |
| Magnification | 105,000x | 105,000x |
| Voltage (kV) | 300 | 300 |
| Electron dose (e^-^/Å) | 60 | 60 |
| Dose per frame (e^-^/Å) | 0.93 | 0.93 |
| Defocus range (μm) | -1.3 to -1.8 | -1.3 to -1.8 |
| Pixel size (Å) | 0.828 | 0.828 |
| Symmetry imposed | C2 | C1 |
| Initial number of particles | 1,573,299 | 1,573,299 |
| Final particle number | 675,686 | 350,283 |
| Map resolution (Å) | 2.23 | 2.54 |
| Map resolution range (Å) | 1.83-3.99 | 1.81-5.57 |
| FSC threshold | 0.143 | 0.143 |
| Refinement |  |  |
| Initial model used | 1IRU | 1IRU |
| Model resolution (Å) | 2.31 | 2.65 |
| FSC threshold | 0.5 | 0.5 |
| Map-sharpening B-factor (Å^2^) | -83.2 | -89.4 |
| Model composition |  |  |
| Non-hydrogen atoms | 48,892 | 49,780 |
| Protein residues | 6,280 | 6,404 |
| *B*-factors (Å^2^) |  |  |
| Protein | 34.37 | 41.91 |
| R.m.s. deviations |  |  |
| Bond lengths (Å) | 0.005 | 0.002 |
| Bond angles (°) | 0.601 | 0.429 |
| Validation |  |  |
| MolProbity score | 1.21 | 1.36 |
| Clashscore | 4.31 | 5.23 |
| Poor rotamers (%) | 0.19 | 1.26 |
| Ramachandran statistics (%) |  |  |
| Favored | 98.60 | 98.60 |
| Allowed | 1.40 | 1.40 |
| utliers | 0 | 0 |
